# Simple contextual cueing prevents retroactive interference in short-term perceptual training of orientation detection tasks

**DOI:** 10.1101/2021.12.15.472760

**Authors:** Hui Huang, Yangming Zhang, Sheng Li

**Affiliations:** School of Psychological and Cognitive Sciences, Peking University, Beijing, China; Beijing Key Laboratory of Behavior and Mental Health, Peking University, Beijing, China; PKU-IDG/McGovern Institute for Brain Research, Peking University, Beijing, China; Key Laboratory of Machine Perception (Ministry of Education), Peking University, Beijing, China

## Abstract

Perceptual training of multiple tasks suffers from interference between the trained tasks. Here, we conducted four psychophysical experiments with separate groups of participants to investigate the possibility of preventing the interference in short-term perceptual training. We trained the participants to detect two orientations of Gabor stimuli in two adjacent days at the same retinal location and examined the interference of training effects between the two orientations. The results showed significant retroactive interference from the second orientation to the first orientation (Experiments 1 and 2). Introducing a 6-hour interval between the pre-test and training of the second orientation did not eliminate the interference effect, excluding the interpretation of disrupted reconsolidation as the pre-test of the second orientation may reactivate and destabilize the representation of the first orientation (Experiment 3). Finally, the training of the two orientations was accompanied by fixations in two colors, each served as a contextual cue for one orientation. The results showed that the retroactive interference was not evident after introducing these passively perceived contextual cues (Experiment 4). Our findings suggest that the retroactive interference effect in short-term perceptual training of orientation detection tasks was likely the result of higher-level factors such as shared contextual cues embedded in the tasks. The effect of multiple perceptual training could be facilitated by associating the trained tasks with different contextual cues.

---

Perceptual training is essential for individuals who need specialized perceptual skills (Deveau et al., 2014; Frank et al., 2020) or suffer perceptual deficits (Huang et al., 2008; Levi & Li, 2009; Polat et al., 2004; Zhang et al., 2014). Previous investigations have shown improved performance for perceptual tasks by training and attributed the training effects to modifications on sensory (Chen et al., 2015; Furmanski et al., 2004; Jia et al., 2020; Schoups et al., 2001; Schwartz et al., 2002; Yan et al., 2014; Yang & Maunsell, 2004; Zivari Adab & Vogels, 2011), decision making (Dosher et al., 2013; Jia et al., 2018; Kahnt et al., 2011; Kuai et al., 2013; Law & Gold, 2008), and short-term memory (Jia et al., 2021; Zhang et al., 2016) processes. A classical finding in perceptual learning was that the training effects were largely specific to the trained tasks and stimuli (Ahissar & Hochstein, 1997; Fahle, 2005; Karni & Sagi, 1991), potentially limiting the applicability of perceptual training outside the laboratories. Transfers of training effects between tasks and stimuli were observed but the generalization of perceptual learning was suggested to be influenced by various factors (Censor, 2013; Jeter et al., 2010; Larcombe et al., 2017; McGovern et al., 2012). Accordingly, previous literatures have proposed different procedures to improve the transfer of training effects to untrained stimuli (Donovan & Carrasco, 2018; Donovan et al., 2015; Wang et al., 2014; Xiao et al., 2008; Zhang et al., 2010). However, given the potential complexity of the tasks and stimuli, the generalization issue remains a barrier for the real-world applications of perceptual training. Alternatively, if the required perceptual skills are within a specific collection of expertise (e.g., for an athlete or a radiologist; Deveau et al., 2014; Evans et al., 2013; Lago et al., 2021), training on a limited number of representative tasks or stimuli may facilitate the wider application of perceptual learning.

Nevertheless, training multiple tasks or stimuli has its own shortcoming as interference is a ubiquitous phenomenon of learning and memory (Anderson, 2003; Anderson et al., 1994; Herszage & Censor, 2018; Wimber et al., 2009). In perceptual learning literature, it has been shown that performance improvement of one task can be disrupted when the initial training is immediately followed by training on a second task (Bang, Shibata, et al., 2018; Been et al., 2011; Seitz et al., 2005; Shibata et al., 2017; Yuko Yotsumoto et al., 2009). This is a typical form of retroactive interference observed in memory research (BrashersKrug et al., 1996; Osgood, 1948; Postman & Underwood, 1973). For example, Seitz and his colleagues have demonstrated that retroactive interference occurs in perceptual learning with a vernier acuity tasks (Seitz et al., 2005). Specifically, in their study, the second task interfered with the training effect of the first task only when the stimuli in the two tasks were in the same orientation and presented at the same retinal location. This finding suggested that the interference of perceptual memory in the vernier task was specific to location and orientation and thus could be attributed to low-level visual processing (e.g., primary visual cortex). The shared neural populations for encoding and storage between the two trained tasks might underlie the observed interference effects.

The classical theory of memory interference highlighted the emerged competition when a retrieval cue is associated with multiple items in the memory (Anderson et al., 1994; Anderson & Neely, 1996; Greeno, 1964; Postman & Underwood, 1973). That is, in memory retrieval, a successful access to a target item from a cue depends not only on how strongly the cue is related to the target item, but also on the strength of the associations between the cue and other distracting items in the memory (Anderson et al., 1994; Anderson & Neely, 1996).

When the cue-target association is not stronger enough than those between the cue and other competing items, memory interference occurs. The phenomenon of interference suggests that the already consolidated memory could suffer from a new learning.

In the present study, we first examined whether the consolidated perceptual memory from a short-term training could be interfered by new perceptual training on the same task. We then investigated the nature of the interference and the potential approach to prevent such interference. We conducted four psychophysical experiments with separate groups of participants (30 participants in each group and 120 participants in total). Figure 1 summarizes the experimental procedures of the four experiments. Experiment 1 established a baseline training effect of an orientation detection task for the other three experiments. In Experiment 2, a retroactive interference effect was observed when two orientations were trained at the same retinal location on two adjacent days. Experiment 3 added a 6-hour interval between the pre-test and training of the second orientation to examine the reactivation account of the interference effect. Finally, Experiment 4 was conducted with two contextual color cues associated with the two orientations when they were trained and tested. This manipulation eliminated the retroactive interference, indicating that higher-level factors related to mnemonic processing may underlie the observed interference effect.

**Figure 1.**
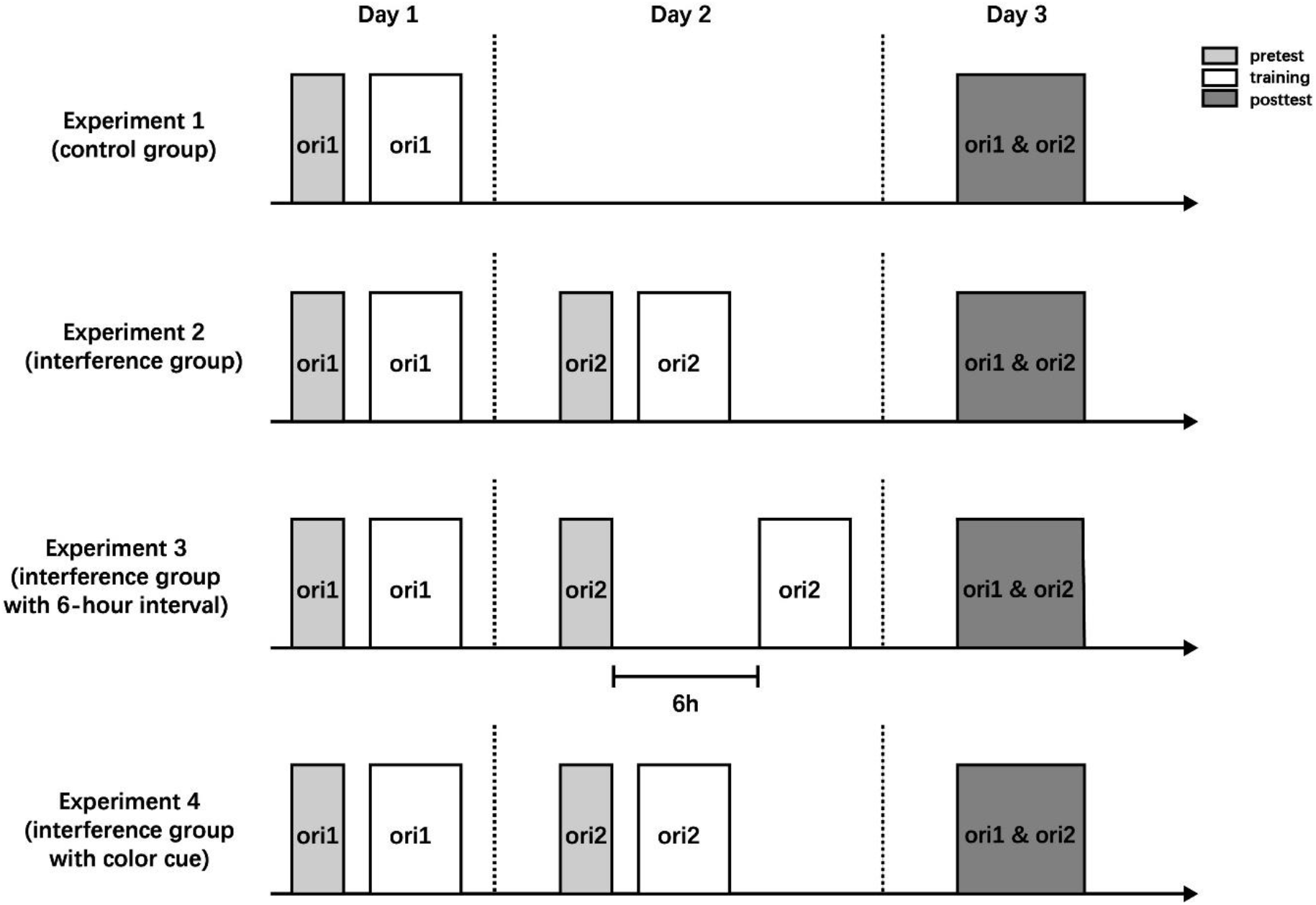
Overview of the procedures for the four experiments.

## Experiment 1: Control group

In Experiment 1, we established the baseline training effect on the orientation detection task when the Gabor stimuli were centrally presented. The participants were trained on one orientation (ori_1) and tested before and after the training. The post-test of ori_1 was conducted at the third day to match with other experiments.

### Method

#### Participants

Thirty right-handed naïve participants (23 females, age range = 18~28years, mean age = 21.47 years) with normal or corrected-to-normal vision were recruited for the experiment. The local ethics committee approved the study.

#### Stimuli and Apparatus

All stimuli were generated in MATLAB (MathWorks, Natick, MA, USA) using Psychtoolbox 3 package (Brainard, 1997; Pelli, 1997) and presented on a gamma-corrected CRT display (1024 × 768 resolution, 85 Hz refresh rate). Subjects viewed the stimuli at a distance of 57 cm with their head stabilized using a chinrest. Gabor patches (spatial frequency = 1 cycle/degree, contrast = 100%, Gaussian filter sigma = 2.5, random spatial phase, radius = 5°) were presented at the center of the display surrounded with gray background (mean luminance = 33 cd/m^2^) during all sessions. A noise pattern created from a sinusoidal luminance distribution was added to the Gabor patches at a given signal-to-noise (S/N). For example, a 10% S/N ratio represented the noise pattern replaced 90% pixels of the Gabor patches.

#### Procedure

##### Orientation detection task

Participants completed a two-interval forced-choice orientation detection task used in previous studies of perceptual learning (Figure 2A, Bang, Sasaki, et al., 2018; Bang, Shibata, et al., 2018; Shibata et al., 2017). Each trial began with a central fixation dot for 500 ms, followed by two 50 ms stimulus intervals. The two stimulus intervals were separated by a 300 ms blank interval with only central fixation displayed. A Gabor patch with a certain S/N ratio and a patch of full noise (0% S/N ratio) were presented in the two stimulus intervals, with their order randomized. The participants were required to determine in which interval the Gabor appeared by pressing one of two buttons on the keyboard. There was no time limit for the response and no feedback was provided.

**Figure 2.**
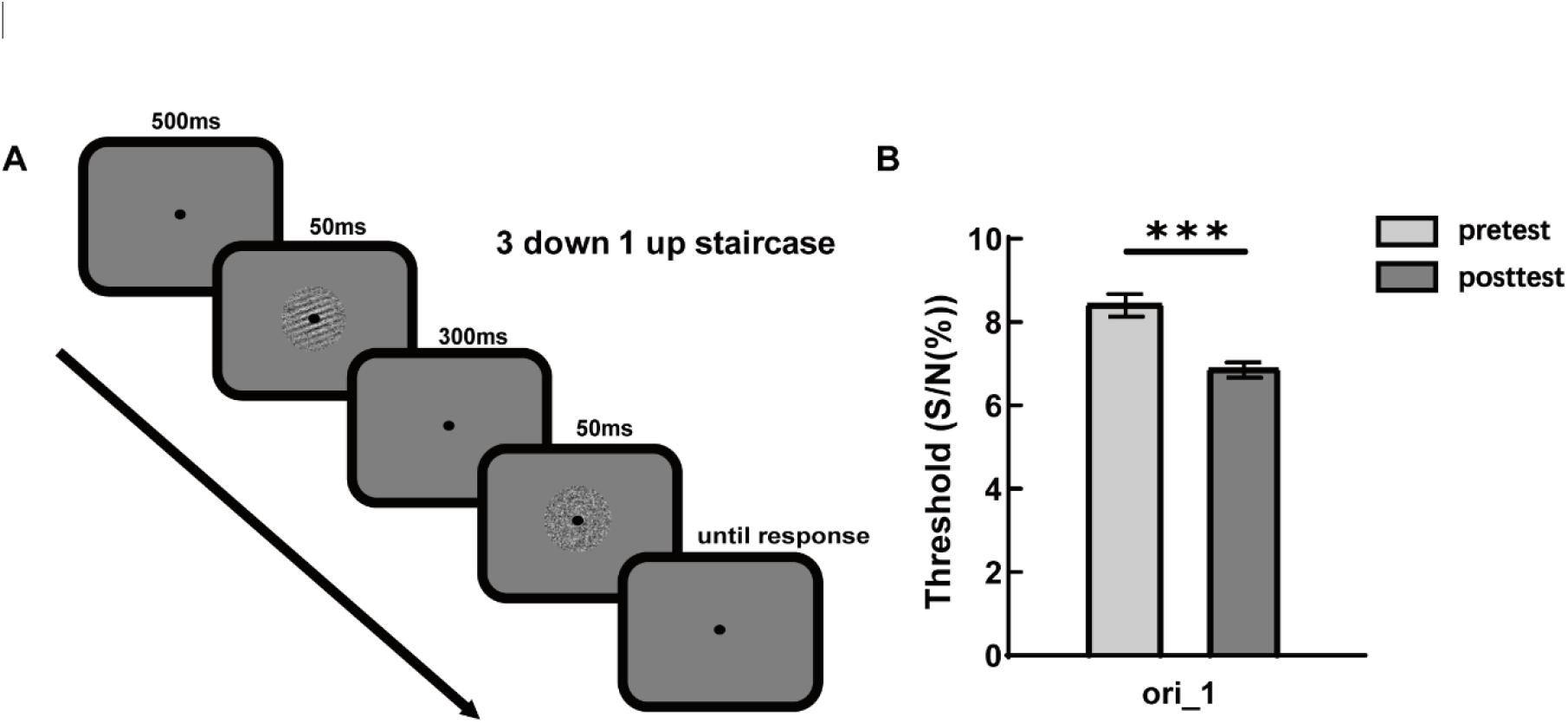
Method and results of Experiment 1. (A) Procedure for an example trial. (B) Behavioral results: thresholds on ori_1 in pre-test and post-test sessions. *** *p* < 0.001. Error bars represent standard errors.

##### Threshold measurement

In the orientation detection task, we used a three-down one-up staircase method to measure each participant’s threshold S/N ratio in each block. This method converged to 79.4% correct responses. In each block, the S/N ratio started with 25% and adaptively changed with a step size of 0.05 log units. Each staircase consisted of four practice and six experimental reversals. The participant’s threshold within a block was defined as the geometric mean of the experimental reversals. The first blocks in the test sessions (including pre-test and post-test) were discarded and we took the arithmetic average of the threshold S/N ratios across the remaining two blocks as the threshold S/N ratio for the tested orientation.

##### Training and test sessions

We adopted two orientations (10° and 70° from the horizontal axis) and they were randomly assigned as ori_1 and ori_2 across participants. The participants in Experiment 1 were regarded as the control group and were trained only on ori_1. On day 1, the participants completed a pre-test session of three blocks on ori_1 and then a training session of 16 blocks on ori_1. There was no task on day 2. On day 3, the participants completed post-test sessions of three blocks on ori_1 and ori_2.

#### Data analysis

To reduce the impact of initial threshold on the training effect, we considered mean percent improvement (MPI) as the dependent variable. MPI depicts the performance improvement and is defined by ((pretest threshold on ori_1 - posttest threshold on ori_1)/pretest threshold on ori_1) ×100 for ori_1. The calculation of MPI for ori_2 is the same as that for ori_1.

### Results and Discussion

As shown in Figure 2B, we observed a significant learning effect on the threshold of ori_1 (t(29) = 7.214, p < 0.001, Cohen’s d = 1.32). The MPI of ori_1 in Experiment 1 served as a baseline training effect for both ori_1 and ori_2 trainings in the following experiments.

## Experiment 2: Interference group

In Experiment 2, after the training of first orientation (ori_1) on day 1, we trained the participants on another orientation (ori_2) on day 2 and examined the training effects of both orientations on day 3. The interferences between the two trainings were measured by comparing their MPIs between Experiment 1 and Experiment 2.

### Method

#### Participants

Thirty right-handed naïve participants (24 females, age range = 18~26 years, mean age = 21.00 years) with normal or corrected-to-normal vision were included in the experiment.

#### Stimuli and Apparatus

The stimuli and apparatus in Experiment 2 were identical to Experiment 1.

#### Procedure

The procedure in Experiment 2 was identical to Experiment 1 with two exceptions. First, the participants completed a pre-test session of three blocks on ori_2 and then a training session of 16 blocks on ori_2 on day 2. Second, the testing order of the two orientations was counterbalanced across participants on day 3.

### Results and Discussion

As shown in Figure 3A, we observed significant learning effects on both ori_1 (t(29) = 3.697, p < 0.001, Cohen’s d = 0.67) and ori_2 (t(29) = 4.449, p < 0.001, Cohen’s d = 0.81).

**Figure 3.**
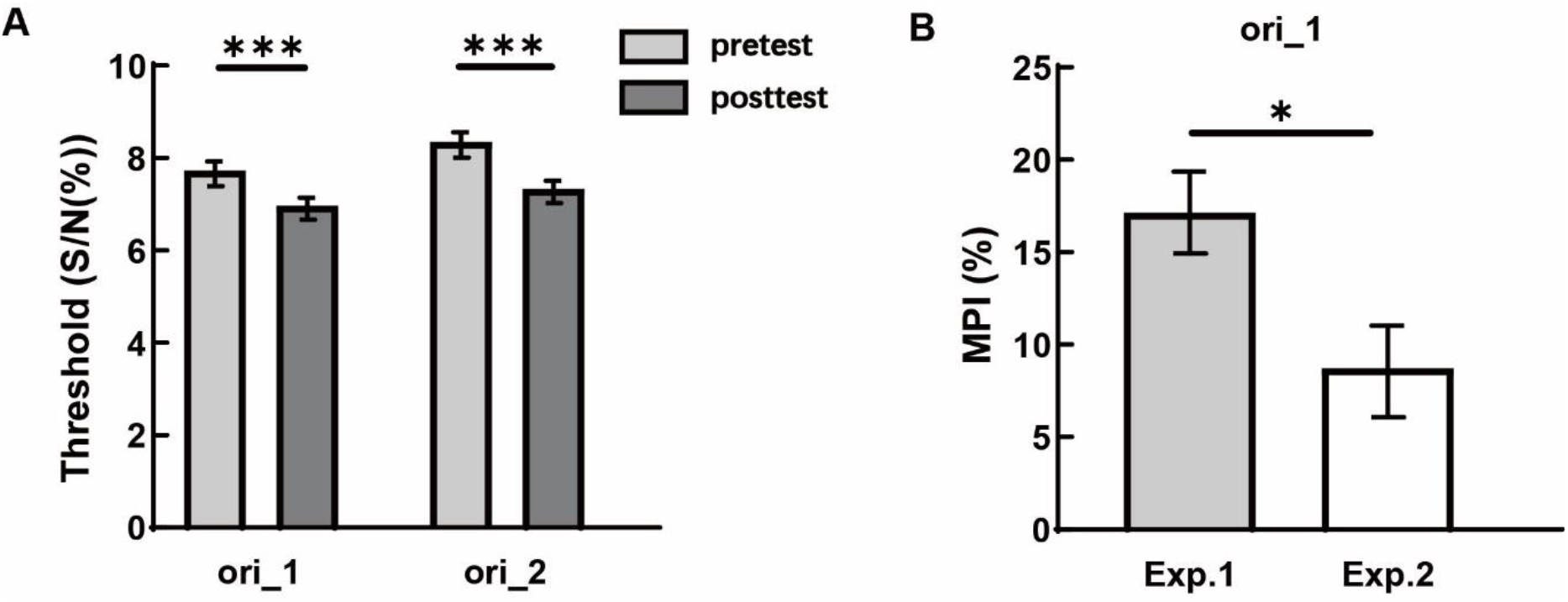
Results of Experiment 2. (A) Behavioral results: thresholds on ori_1 and ori_2 in pre-test and post-test sessions. (B) MPI comparison for ori_1 between Experiment 1 and Experiment 2. * *p* < 0.05, *** *p* < 0.001. Error bars represent standard errors.

To examine whether training on ori_2 would introduce retroactive interference on the learning effect of ori_1, we compared the MPIs of ori_1 between Experiment 1 and Experiment 2. The result revealed that the MPI of Experiment 2 was significantly smaller than that of Experiment 1 (t(58) = 2.528, p = 0.014, Cohen’s d = 0.65, Figure 3B). These results suggest that training on the detection of the second orientation induced retroactive interference on the previously trained detection of the first orientation. To test the possible proactive interference from ori_1 training to ori_2 training, we compared the MPI of ori_2 in Experiment 2 with the MPI of ori_1 in Experiment 1. The result revealed a trend of significant difference between the two MPIs (t(58) = 1.782, p = 0.08, Cohen’s d = 0.46).

We considered a few factors that may drive the observed retroactive interference effect. We first considered the reactivation account. The stimuli of ori_1 and ori_2 were similar in their shapes and other features (e.g., they were both Gabor patches). The training procedures were also identical for the two orientations. Therefore, the learning traces of the two trainings could be activated by each other during their retrievals. In this case, it was possible that pre-test of ori_2 may have reactivated the consolidated learning trace of ori_1 and brought it to an unstable status that was susceptible to interruption (Bang, Shibata, et al., 2018; Censor et al., 2010; Nader, 2015; Schiller et al., 2010). Once destabilized, the old learning trace of ori_1 could be disrupted by the following training on ori_2 and the retroactive interference would be observed. To address this possible interpretation, we conducted Experiment 3 in which a 6-hour interval was introduced between the pre-test and training of ori_2. The length of the interval was determined by previous literatures showing that the reactivated memory becomes reconsolidated 6 hours after reactivation (Bang, Shibata, et al., 2018; Censor et al., 2010; Huang & Li, 2021; Nader, 2015; Schiller et al., 2010; Shibata et al., 2017). If the reactivation account could explain the observed retroactive interference, we would expect to observe eliminated interference effect in Experiment 3.

## Experiment 3: Interference group with a 6-hour interval

### Method

#### Participants

Thirty right-handed naïve participants (22 females, age range = 18~27 years, mean age = 22.23 years) with normal or corrected-to-normal vision were included in the experiment.

#### Stimuli and Apparatus

The stimuli and apparatus in Experiment 3 were identical to Experiment 1.

#### Procedure

The general procedure in Experiment 3 was identical to Experiment 2, except that on day 2, there was a 6-hour interval between the pre-test and training sessions of ori_2.

### Results and Discussion

As shown in Figure 4A, we observed significant learning effects on both ori_1 (t(29) = 2.141, p = 0.041, Cohen’s d = 0.39) and ori_2 (t(29) = 5.511, p < 0.001, Cohen’s d = 1.01).

**Figure 4.**
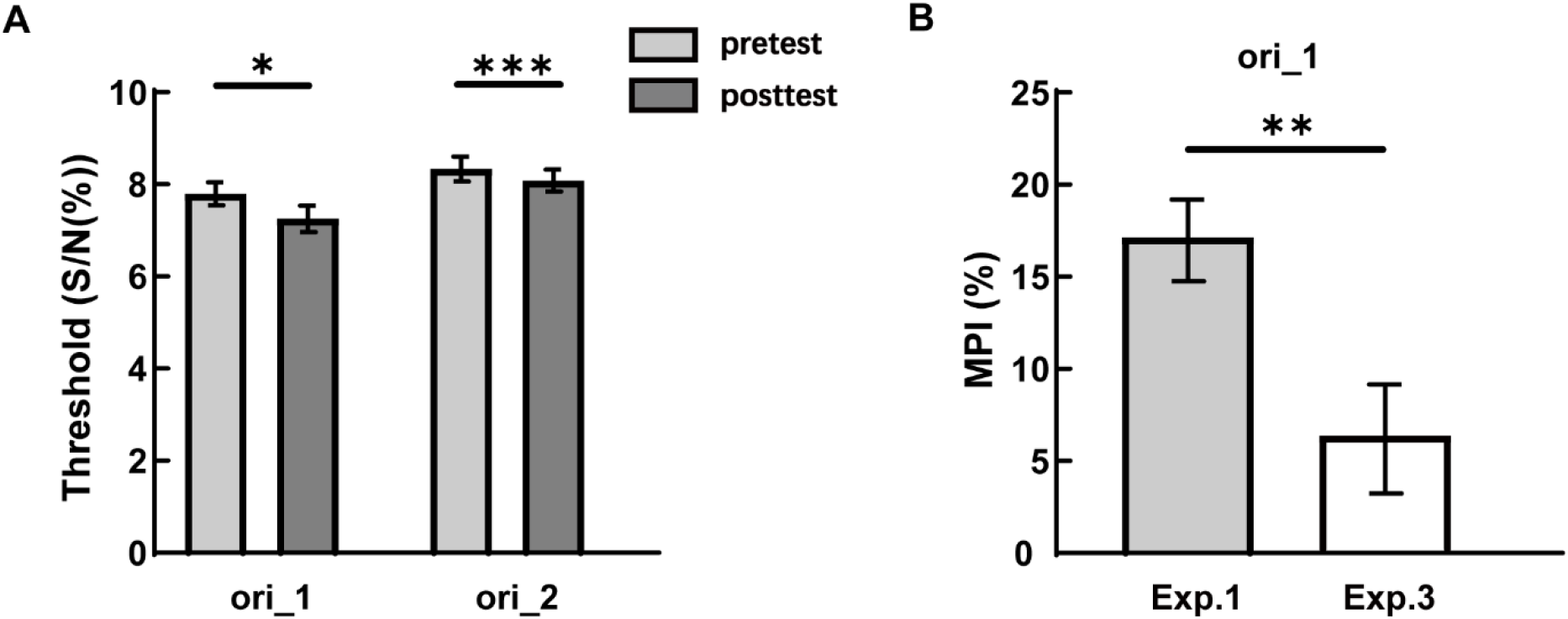
Results of Experiment 3. (A) Behavioral results: thresholds on ori_1 and ori_2 in pre-test and post-test sessions. (B) MPI comparison for ori_1 between Experiment 1 and Experiment 3. * *p* < 0.05, ** *p* < 0.01, *** *p* < 0.001. Error bars represent standard errors.

We compared the MPIs of ori_1 between Experiment 1 and Experiment 3 and the result revealed that the MPI of Experiment 3 was significantly smaller than that of Experiment 1 (t(58) = 2.956, p = 0.005, Cohen’s d = 0.55, Figure 4B). We also compared the MPI of ori_2 in Experiment 3 with the MPI of ori_1 in Experiment 1 and the result revealed no significant difference between the two MPIs (t(58) = 0.754, p = 0.454, Cohen’s d = 0.19). These results demonstrated a similar retroactive interference effect as in Experiment 2 when a six-hour interval was introduced between the pre-test and training of ori_2. The results suggest that the observed retroactive interference effect could not be explained by the destabilized status of the learning trace of ori_1 due to the pre-test of ori_2. Therefore, the reactivation account was unlikely a sufficient interpretation.

There were two other accounts that could serve as the potential interpretations for the observed interference effect. First, training to detect the Gabor stimuli recruited orientation selective neurons at the primary visual cortex. The neurons involved in the training may not be restricted to those selective to the trained orientation as only a fraction of neurons in visual cortex are dedicated to encode orientation information (Poort et al., 2015). However, the neurons recruited by both ori_1 and ori_2 training may come from an overlapping population of neurons that were retinotopically responded to the same stimulus location. The interference could occur from the new training to the old training as they would compete for the recruitment of the same population of neurons. We referred this possibility as the competing neuronal population account. Second, training may involve the establishment of association between stimuli’s orientations and training contexts that could benefit the retrieval of the learning traces during test. If, however, the two orientations were linked to the same contextual cue (e.g., training protocol or task set), retrieval errors would occur during the test session. The retrieval failure due to shared contextual cue could also contribute to the observed interference effect. We referred this possibility as the shared context account.

We conducted Experiment 4 to dissociate these two possible interpretations. In Experiment 4, the two orientations were associated with different colors during training and tests. If the competing neuronal population account was true, we would again observe retroactive interference. If the shared context account was true, associating two orientations with two colors would prevent the retroactive interference effect.

## Experiment 4: Interference group with color cues

### Method

#### Participants

Thirty right-handed naïve participants (16 females, age range = 18~29 years, mean age = 21.67 years) with normal or corrected-to-normal vision were included in the experiment.

#### Stimuli and Apparatus

The stimuli and apparatus in Experiment 4 were identical to Experiment 1.

#### Procedure

The general procedure in Experiment 4 was identical to Experiment 2, except that the Gabor stimuli of ori_1 and ori_2 were associated with fixation of different colors (red or green) in the training and test sessions (Figure 5A). The color-orientation associations were counterbalanced across participants. The participants received no instruction about the fixation colors.

**Figure 5.**
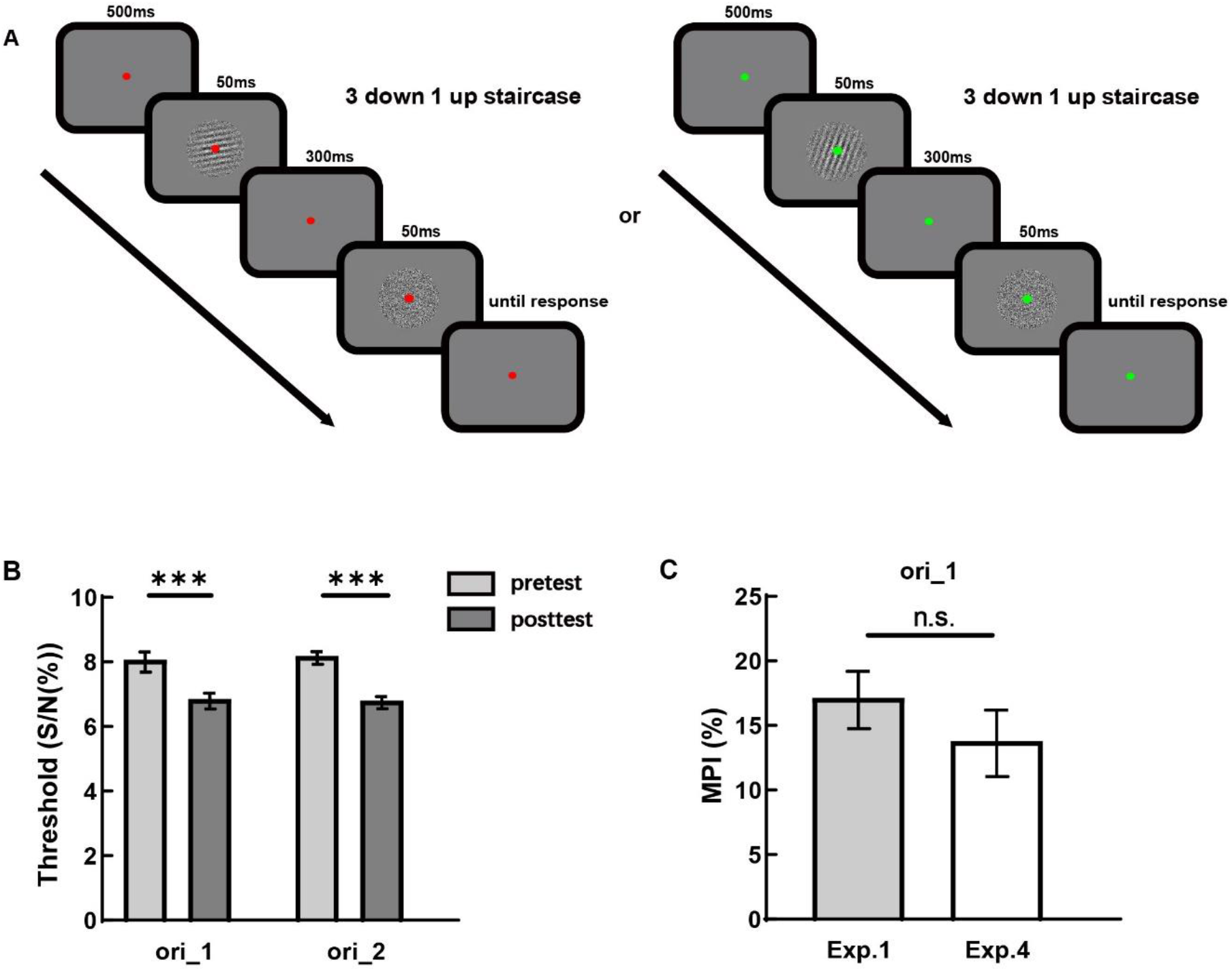
Method and results of Experiment 4. (A) Procedure for an example trial. The two orientations were associated with two colors (red or green) during the training and test sessions. (B) Behavioral results: thresholds on ori_1 and ori_2 in pre-test and post-test sessions. (C) MPI comparison for ori_1 between Experiment 1 and Experiment 4. *** *p* < 0.001. Error bars represent standard errors.

### Results and Discussion

As shown in Figure 5B, we observed significant learning effects on both ori_1 (t(29) = 5.154, p < 0.001, Cohen’s d = 0.94) and ori_2 (t(29) = 6.941, p < 0.001, Cohen’s d = 1.27).

We compared the MPIs of ori_1 between Experiment 1 and Experiment 4 and the result revealed that the MPI of Experiment 4 was not significantly different from that of Experiment 1 (t(58) = 0.986, p = 0.328, Cohen’s d = 0.25, Figure 5C), suggesting that the first training would not be interfered by the second training if the two orientations were associated with different color cues. No significant difference between the MPI of ori_2 in Experiment 4 with the MPI of ori_1 in Experiment 1 was observed (t(58) = 0.284, p = 0.777, Cohen’s d = 0.07).

These results were not consistent with the competing neuronal population account and provided supporting evidence for the shared context account. That is, the shared contextual cue between the two orientations introduced the retroactive interference in Experiment 2. The results demonstrated that relating the two orientations with different contextual cues could prevent the retroactive interference effect.

## General Discussion

The present study investigated the interference effects between two orientations that were trained in the same detection task in two adjacent days. The results showed that training on the second orientation impaired the training effect of the first orientation, demonstrating a retroactive interference effect (Experiment 1 and Experiment 2). This effect could not be explained by the reactivation account and competing neuronal population account, as were evident in Experiment 3 and Experiment 4, respectively. Finally, associating the two orientations with two contextual color cues during the training and test prevented the emergence of the retroactive interference effect, providing the supporting evidence to the shared context account (Experiment 4). These findings suggest that multiple short-term perceptual training could suffer from retroactive interference which is likely the result of higher-level factors such as shared contextual cues embedded in the tasks. To facilitate the efficiency of multiple training, associating the trained tasks with different contextual cues could be a solution in practice.

Perceptual learning has been argued to arise from plastic changes in primary sensory areas (Ahissar & Hochstein, 1997; Bao et al., 2010; Jehee et al., 2012; Schoups et al., 2001; Schoups et al., 1995; Schwartz et al., 2002; Yan et al., 2014). The training protocol adopted in the present study, despite short in time, nevertheless resulted in behavioral improvement on the trained task after one-day consolidation. The retroactive interference effect observed in Experiment 2 where the two orientations were trained at the same retinal location could be attributed to a shared pool of neurons in primary visual cortex that responded to both orientations (Seitz et al., 2005). However, this interpretation was not supported by the absent of retroactive interference effect in Experiment 4 in which the two orientations were in the same retinal location but associated with different colors during the training and test. These results supported another interpretation that the shared contextual cues between the two orientations might be a critical factor.

Nevertheless, our conclusion did not imply that no plastic change occurred in the primary visual cortex. In fact, previous studies have demonstrated that, using perceptual training paradigm with Gabor stimuli, training effect transferred little to a new orientation if it was at the same retinal location as the trained orientation (Dosher et al., 2013). The orientation specificity of the training effect suggests that training of the two orientations may not recruit the overlapping population of neurons in the primary visual cortex, further excluding the competing neuronal population account for the interpretation of our results. It was more likely that training of the two orientations in Experiment 2 involved two populations of orientation-selective neurons at the primary visual cortex but established shared associations between the orientations and contextual cues (e.g., training protocol or task set). If performing the trained task initiated a retrieval process that depended on the orientation-context association, the retrieval of the perceptual memory would have been disrupted by the shared context between the two orientations. By introducing orientation-color associations in Experiment 4, the two orientations were associated with different contextual cues and the reliable retrieval of their learning traces became possible. The recovery from retroactive interference by the orientation-color associations provided us with new insight about the retrieval of learned perceptual skills. In Experiment 4, the participants were not informed about the meaning of the color cues. They just passively perceived the cues during the training and test. Our results suggest that the contextual cue could serve as a critical factor for the successful retrieval of multiple perceptual learning traces. This could be achieved by facilitating hippocampus-dependent pattern separation during the training phase as the hippocampus is known to play an important role in forming distinct memory traces for similar experiences (O’Reilly & Rudy, 2001). We suggest that future investigations with neuroscientific approaches are required for revealing a full picture of the interaction between memory retrieval and sensory plasticity in perceptual learning.

There were a few studies in the literature that investigated the interferences between multiple perceptual trainings. In most of these studies, interferences were observed when the second training were immediately after or separated up to a few hours from the first training (Been et al., 2011; Seitz et al., 2005; Shibata et al., 2017; Yuko Yotsumoto et al., 2009). That is, the two trainings were conducted at the same day and the within-day switchover of the two trainings lasted for several days. Seitz et al. (2005) has demonstrated with a vernier task that multiple perceptual training could induce feature- and location-specific interference when the second training started immediately after the first training. However, the interference was not evident if the two trainings were separated by one hour, suggesting well-consolidated memory could prevent the interference from later training. In another study that adopted Gabor stimuli with an orientation discrimination task, Been et al. (2011) also revealed retroactive interferences in conditions in which the intervals between the two trainings varied from 0 to 24 hours. Their results suggested that the overlap between neuronal populations of the two trained orientations was responsible for the interference. No evidence for the role of memory consolidation was revealed in this study.

In contrast to the findings in Seitz et al. (2005) and Been et al. (2011), we did not find evidence for the perceptual interpretation of the interference effects. There are three possible reasons for the different findings. First, as mentioned above, the first orientation was well-consolidated with an overnight sleep before the second training in the present study, whereas the two trainings were separated up to a few hours in these two studies (except the 24-hour condition in Been et al. 2011). Sleep-related consolidation was known to modulate perceptual learning (Tamaki et al., 2020; Yotsumoto et al., 2009). It is reasonable to consider the possibility that sleep also influences the interference mechanism of perceptual memory. Second, an outstanding difference between our study and these two studies was the task used for training. Compared with the orientation detection task, vernier task and orientation discrimination task were considered relying more on primary visual cortex (Crist et al., 1997; Fahle, 1997; Fiorentini & Berardi, 1981; Shiu & Pashler, 1992). The difference in task could contribute to different levels of competition between neuronal populations recruited by two trainings. Third, we adopted a short training approach in which each orientation was trained for one day, whereas each training was repeated for five (Seitz et al., 2005) and fifteen (Been et al., 2011) days in these two studies. It was known that extensive training resulted in precise discrimination and involves lower-level processing (Ahissar & Hochstein, 1997; Hochstein & Ahissar, 2002). Thus, it is reasonable to suggest that the level at which the interference occurred was related to the level of training-related processing. As a result, we suggest that the interference induced by multiple perceptual training could be modulated by low-level perceptual processing or higher-level mnemonic mechanisms, given the different tasks to be trained.

In a study that investigated the reconsolidation effect in perceptual learning, Bang et al. (2018) adopted a Gabor orientation detection task as in the present study. Particularly, the long interval condition in their study 1 was similar to our Experiment 2 except that there was a test for the first orientation 3.5 hours before the pre-test of the second orientation in day 2. However, unlike our results that showed significant retroactive interference effect, their results revealed intact training effect for the first orientation in day 3 (in their Fig. 1c). As suggested by their findings, the test of the first orientation in day 2 induced reactivation and reconsolidation processes. It has long been known that retrieval practice (i.e., testing) could facilitate the long-term retention of memory(Roediger & Karpicke, 2006). Therefore, the test of the first orientation in day 2 in their study would strengthen its original training effect. As suggested by another study of the same group (Shibata et al., 2017), the strengthened learning could stabilize perceptual training effect and protect it from retroactive interference of a new learning (also see (Potts & Shanks, 2012)for the case in associative learning). In our Experiment 2, there was no test for ori_1 in day 2 and this may explain the mentioned difference between the two studies. Future investigations are required to further elucidate the relationship between memory reconsolidation and perceptual training interference.

We also examined the possible proactive interference from the training of the first orientation to the training of the second orientation. We compared the MPIs of the second orientation in Experiments 2, 3, and 4 with the MPI in Experiment 1. Apart from a trend of significance in comparing Experiment 2 and Experiment 1, we did not find other significant difference. We noted that the post-test in Experiment 1 was two days after the pre-test and training and this gap in the other three experiments was one day. Previous study has suggested that proactive interference was more likely to happen if there was overlearning for the first trained task (Shibata et al., 2017). In the present study, two trainings were both short-term and equalized in terms of training length. Therefore, we suggest that there was only weak, if not absent, proactive interference the present study.

Roving as a training method for perceptual learning was extensively studied in the literature (Cong & Zhang, 2014; Dosher et al., 2020; Kuai et al., 2005; Parkosadze et al., 2008; Tartaglia et al., 2009; Yu et al., 2004; Zhang et al., 2008). Different from the present study, in a roving experiment, trials of different conditions were randomly interleaved during the training, preventing subjects from learning the trained feature due to the interference between trials. Interestingly, Zhang et al. (2008) conducted a contrast discrimination experiment in which four roving contrasts (i.e., four stimulus conditions) were each assigned a tag of letter. They found that the tagging prevented learning from the disruption of roving and suggested that semantic tags could facilitate top-down attention during training. The contextual color cueing in our Experiment 4 shared a similar associative idea with the tagging manipulation in Zhang et al. (2008), but focused more on the retrieval process underlying the interference effects in perceptual training. The results of both studies suggested the importance of mnemonic mechanisms in understanding perceptual training.

In summary, we demonstrated that two short-term perceptual trainings suffered from retroactive interference due to shared context related to training protocol or task set. Introducing color-based contextual cues could establish separate contexts for the two trainings and prevent the occurrence of retroactive interference. Hippocampus-dependent pattern separation was likely to play a critical role in this process but this idea needs future neuroscience investigations to confirm. Overall, our findings emphasized the roles of mnemonic mechanisms in fully understanding the phenomenon of perceptual learning.

